# Forward genetics combined with unsupervised classifications identified zebrafish mutants affecting biliary system formation

**DOI:** 10.1101/2021.06.22.449425

**Authors:** Divya Jyoti Singh, Kathryn M. Tuscano, Karen L. Ortega, Manali Dimri, Kevin Tae, William Lee, Muslim A. Muslim, Jay L. Liu, Lain X. Pierce, Allyson McClendon, Gregory Naegele, Isabel Gibson, Jodi Livesay, Takuya F. Sakaguchi

## Abstract

Impaired formation of the biliary network can lead to congenital cholestatic liver diseases; however, the genes responsible for proper biliary system formation and maintenance have not been fully identified. Combining computational network structure analysis algorithms with a zebrafish forward genetic screen, we identified 24 new zebrafish mutants that display impaired intrahepatic biliary network formation. Complementation tests suggested that these 24 mutants affect 24 different genes. We applied unsupervised clustering algorithms to classify the recovered mutants into three classes unbiasedly. Further computational analyses revealed that each of the recovered mutations in these three classes shows a unique effect on node subtype composition and connection property distribution of the intrahepatic biliary network. Besides, we found that most recovered mutations are viable. In those mutant fish, biliary network phenotypes persist into adulthood, which themselves are good animal models to study chronic cholestatic liver diseases. Altogether, this study provides unique genetic and computational toolsets that advance our understanding of the molecular pathways leading to biliary system malformation and cholestatic liver diseases.

## Introduction

Cholestatic liver diseases are characterized by slowed or blocked bile flow within the liver and are found in patients of almost any age (1, 2). Biliary atresia, a cholestatic liver disease affecting neonatal patients, occurs in approximately one in 10,000 live births (3, 4) and is characterized by blockage, underdevelopment, and malformation of the biliary ducts that leads to buildup of bile and, ultimately, scarring and cirrhosis of the liver. Biliary atresia continues to be the most common indication for liver transplantation and accounts for almost 50% of all liver transplantation performed in children (4). In general, cholestatic liver diseases are caused by a disequilibrium between the loss of biliary epithelial cells (BECs, also called cholangiocytes) and generation of new BECs, thus liver cholestasis could be induced by a variety of upstream signaling pathways such as activation of death receptors, immune-mediated injury, or oxidative stress (1). Identifying signaling pathways that may induce biliary disease is an important step in understanding these diseases, but there is still much that is unknown. Indeed, relatively few genes that regulate biliary system morphogenesis have been found (5, 6); thus, forward genetic screens could find more genes responsible for this process. Identifying new genes will lead to the discovery of genes susceptible to cholestatic liver diseases and provide new animal models for such diseases.

Systematic forward genetic screens have led to the identification and characterization of genetic pathways that regulate many biological processes, including morphogenesis (7–10). Historically, classifications of mutant phenotypes collected through genetic screens were decided subjectively in which observers decided on classification criteria based on biological knowledge. In contrast, machine learning-based unsupervised classification is an objective process of grouping in which a dataset is sorted based on similarity without any ground truth or pre-labeling of data (11). Unsupervised learning approaches are widely used to make partitioning observations into homogenous clusters as well as to uncover subpopulation structure of hidden relationships in a given dataset (12, 13). This approach has proven to be a powerful tool for exploring genomics and other omics high-dimensional data sets, yet its usefulness for unbiased mutant phenotype characterizations has not been intensively tested.

The zebrafish transgenic line *Tg(Tp1-MmHbb:EGFP)^um14^*, which is generated by conjugating tandem *RBP-Jk* response element repeats with enhanced green fluorescent protein (EGFP), expresses EGFP specifically in the intrahepatic biliary network in the zebrafish liver (14, 15). This transgenic line allowed us to visualize the entire intrahepatic biliary network in zebrafish larvae, greatly facilitating investigation of the development of the biliary system. Utilizing this transgenic line, we have previously developed computational algorithms to convert complex three-dimensional network structures into arrays of numeric sub-parameters to quantify the subtle differences in intrahepatic biliary network branching patterns (16). Precise computational quantification of the intrahepatic biliary network enabled us to unbiasedly quantify mutant phenotypes and break down complicated three-dimensional branching patterns into multi-dimensional arrays, which we can feed into unsupervised classification algorithms.

In this study, we have conducted a forward genetic screen for new mutants that show specific phenotypes in the intrahepatic biliary network, and we recovered 24 of these mutants. We have applied computational network analysis alongside unsupervised classification algorithms to elucidate phenotype groups and unbiasedly revealed new classes of mutant phenotypes. Together, these studies lay the foundation for integrating unsupervised approaches with genetic screening and provide new biological resources to study biliary system biology.

## Results

### A forward genetic screen identified 24 mutants affecting intrahepatic biliary network formation

We have previously developed a computational method to precisely quantify network structural properties of the zebrafish intrahepatic biliary network (16). This method allowed us to characterize subtle differences in network branching and connection patterns. Here, we applied this method to facilitate a forward genetic screen. We completed a forward genetic screen to identify mutants showing altered *Tg(Tp1-MmHbb:EGFP)^um14^* expression in the intrahepatic biliary network at 5 days post-fertilization (dpf) (Materials and Methods). During the screening and recovery procedure, we frequently used the computational skeletal analysis algorithm (16) to confirm that the subtle branching differences we observed in the intrahepatic biliary network of mutant larvae were significantly different from those in wild-type larvae. Since we wanted to identify mutations that specifically affect biliary system morphology, we have excluded any mutations that induced general body shape change, obvious pigmentation change, altered morphology of observable organs, defects in blood circulation, or cardiac edema. Although we observed some interesting biliary system phenotypes in mutants with such observable morphological changes during the screening, we excluded them since the mutations causing such phenotypes were previously screened intensively (17, 18). In the F3 generation, we identified 28 mutations that fit our screening criteria and eventually recovered 24 mutations in the next generation (Table 1). These 24 mutants show consistent phenotypes even after nine or more backcrosses to the original wild-type strain. After we outcrossed the recovered mutants to the wild-type strain for at least five generations, we initiated precise characterization of mutant phenotypes in the intrahepatic biliary network by applying the computational skeletal analysis algorithm (16) (Fig. 1) (Materials and Methods). We converted branching patterns in the *Tg(Tp1-MmHbb:EGFP)^um14^* – expressing intrahepatic biliary network to computational skeletal representation images and corresponding numeric array datasets. We found that these 24 mutants show unique branching patterns in the intrahepatic biliary network that are different from those of the wild-type larvae at 5 dpf (Fig. 1). We did not observe any overt difference in physical appearance among those 24 recovered mutant larvae at 5 dpf, suggesting that those mutations specifically affect the biliary system. Thus, we established 24 mutant alleles responsible for unique intrahepatic biliary network branching phenotypes.

**Table 1.** Mutants recovered from the forward genetic screen.

| **Allele** | **Screen ID** | **Mutant Class** | **Mutation Viability** | **BEC Segregation** | **Sample number** | **Adult Phenotype** |
| --- | --- | --- | --- | --- | --- | --- |
| *Iri11* | AK-13.4 | Class Ia | Viable |  | 6 |  |
| *Iri12* | AK-8.5 | Class Ib | Viable |  | 5 |  |
| *Iri13* | BG-1.2 | Class II | Viable |  | 7 | Slightly reduced |
| *Iri14* | BJ-1.2 | Class Ia | Viable |  | 9 | Slightly reduced |
| *Iri15* | BJ-2.5 | Class II | Viable |  | 9 |  |
| *Iri16* | BJ-3.7 | Class Ia | Lethal | Yes | 8 |  |
| *Iri17* | CD-1.4 | Class Ib | Lethal | Yes | 7 |  |
| *Iri18* | DK-3.2 | Class Ib | Viable |  | 7 |  |
| *Iri19* | DK-3.4 | Class II | Viable |  | 5 |  |
| *Iri20* | EJ-3.10 | Class III | Viable |  | 7 | Thickened |
| *Iri21* | FG-9.3 | Class Ib | Viable |  | 5 | Greatly reduced |
| *Iri24* | GK-1.4 | Class Ia | Viable |  | 5 | Greatly reduced |
| *Iri25* | HK-1.7 | Class Ia | Viable |  | 7 |  |
| *Iri26* | HZ-1.4 | Class Ib | Viable | Yes | 10 | Slightly reduced |
| *Iri27* | IK-1.6 | Class III | Viable |  | 15 |  |
| *Iri29* | IL-2.3 | Class Ib | Viable |  | 9 | Thickened |
| *Iri30* | IL-4.1 | Class Ib | Viable |  | 6 |  |
| *Iri31* | IZ-1.6 | Class Ib | Viable | Yes | 5 | Greatly reduced |
| *Iri33* | JL-7.2 | Class II | Lethal |  | 5 |  |
| *Iri34* | JL-9.3 | Class II | Viable |  | 10 |  |
| *Iri35* | JW-1.10 | Class Ib | Viable |  | 25 |  |
| *Iri36* | JX-2.14 | Class Ia | Lethal |  | 8 |  |
| *Iri37* | KL-10.6 | Class Ib | Viable |  | 10 | Slightly reduced |
| *Iri38* | KZ-3.1 | Class III | Viable |  | 12 |  |

**Figure 1.**
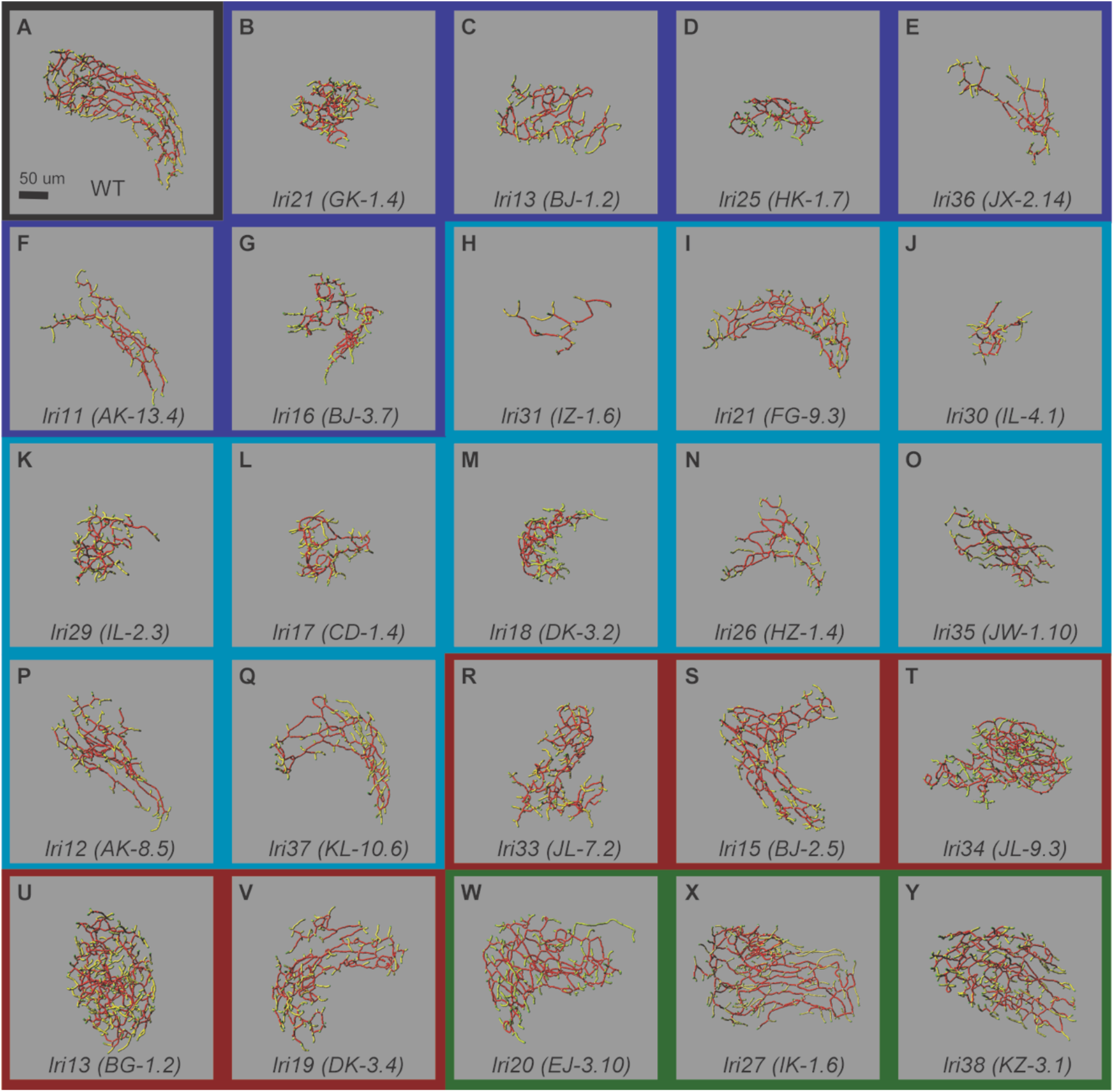
Skeletal representation of the intrahepatic biliary network at 5 dpf in mutants recovered from the forward genetic screen. Skeletal representation of the intrahepatic biliary network in mutant larvae based on *Tg(Tp1-MmHbb:EGFP)^um14^* expression at 5 dpf, unbiasedly classified based on branching sub-parameters of the intrahepatic biliary network. (**A**) Wild-type control (Black border). **(B-G)** Class Ia (Dark Blue border): *lri24* (*GK-1.4*) mutant (B), *lri14* (*BJ-1.2*) (C), *lri25* (*HK-1.7*) (D), *lri36* (*JX-2.14*) (E), *lri11* (*AK-13.4*) (F), and *lri16* (*BJ-3.7*) (G) mutants belong to this class. **(H-Q)** Class Ib (Light Blue border): *lri31* (*IZ-1.6*) (H), *lri21* (*FG-9.3*) (I), *lri30* (*IL-4.1*) (J), *lri29* (*IL-2.3*) (K), *lri17* (*CD-1.4*) (L), *lri18* (*DK-3.2*) (M), *lri26* (*HZ-1.4*) (N), *lri35* (*JW-1.10*) (O), *lri12* (*AK-8.5*) (P), and *lri37* (*KL-10.6*) (Q) mutants belong to this class. **(R-V)** Class II (Red border): *lri33* (*JL-7.2*) (R), *lri15* (*BJ-2.5*) (S), *lri34* (*JL-9.3*) (T), *lri13* (*BG-1.2*) (U), and *lri19* (*DK-3.4*) (V) mutants belong to this class. **(W-Y)** Class III (Green border): *lri20* (*EJ-3.10*) (W), *lri27* (*IK-1.6*) (X), and *lri38* (*KZ-3.1*) (Y) mutants belong to this class. Ventral views, anterior to the top. The network features are color-coded: end points (green), nodes (white), node-node connections (red), and node-end point connections (yellow). The representative biliary skeleton for each genotype was selected based on the lowest deviation from the average network structural sub-parameter attributes.

### Classification of recovered mutants based on structural sub-parameters of the intrahepatic biliary network

Since the skeletal network analysis produces high-dimensional numeric array data of the intrahepatic biliary network in individually examined larva, we applied the hierarchical clustering heat map algorithm (12) to analyze this data. For each genotype, we first calculated the mean values of each sub-parameter obtained from the initial skeletal conversion. We then used the mean values to apply the clustered heat map algorithm to generate the heat map (Fig. 2) showing the phenotypical similarities between all mutants recovered. This is a completely unbiased classification based solely on the structural sub-parameters of the intrahepatic biliary network. Based on the dendrogram of the clustering heat map data (Fig. 2), we assigned three major mutant phenotype classes: Class I, Class II, and Class III. We found that mutants belonging to Class I show decreased total number of segments within the intrahepatic biliary network, indicating that this structural sub-parameter is important for the classification of this group. Based on the dendrogram, this class can be further divided into Class Ia and Class Ib, in which mean network density is increased or decreased, respectively. In both Class II and III, the mean network density is increased compared to that in wild-type larvae, while the total segment number is relatively consistent in Class II but is increased in Class III. Together, these findings suggest that, among all network structural sub-parameters, the total segment number and mean density are relatively more important features for classifying recovered mutant phenotypes.

**Figure 2.**
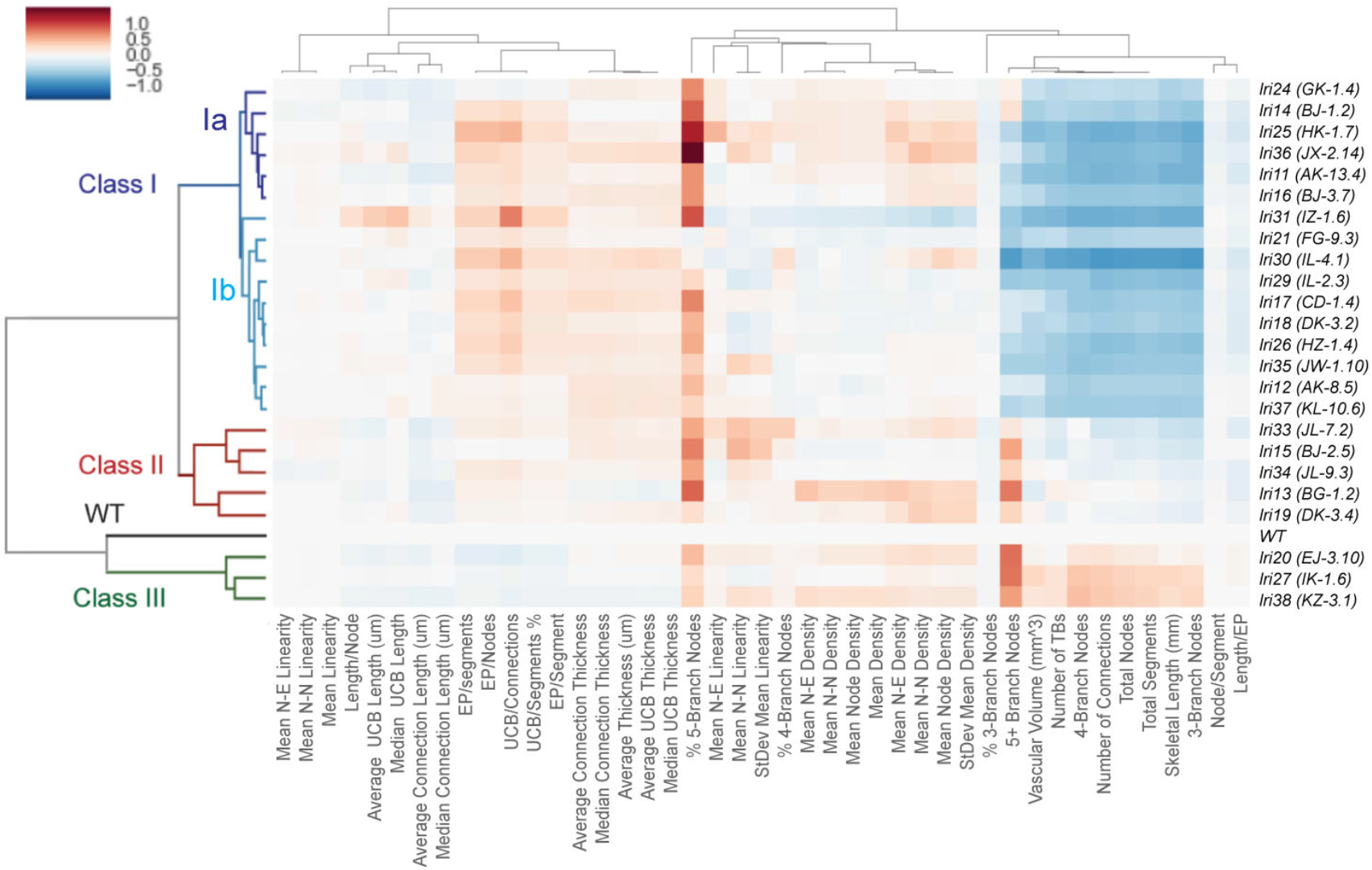
Clustered heat map-based classification of recovered mutants. Clustered heat map based on the structural sub-parameters of the intrahepatic biliary network in recovered mutant larvae at 5 dpf. All sub-parameters are normalized to the average value of the corresponding sub-parameters in the wild-type dataset. Clustering was done with the hierarchical clustering algorithm. Red indicates a higher value than wild-type, and blue indicates a lower value than wild-type. Values represent the average for each specimen dataset. The x-axis of the dendrogram indicates the relative similarity of each structural sub-parameter to other sub-parameters. The shorter the line between two sub-parameters, the more closely related they are. Similarly, the y-axis indicates the relative similarity of each mutant phenotype to other mutant phenotypes. Classes are denoted by the color of the y-axis dendrogram: Wild-type (Black), Class Ia (Dark Blue), Class Ib (Light Blue), Class II (Red), and Class III (Green). Classification was unbiasedly based on network structural sub-parameters of the intrahepatic biliary network.

However, it is possible that the use of clustered heat map analysis does not always respect the intrinsic relationships of mutant phenotypes. In order to independently confirm the phenotypical relationships, we have applied principal component analysis (PCA) (12), which is a different dimensionality-reduction algorithm (Fig. 3). We found that the mutant phenotype classes assigned based on the clustered heat map analysis are grouped together in the plot, suggesting that both algorithms similarly classified the phenotypes.

**Figure 3.**
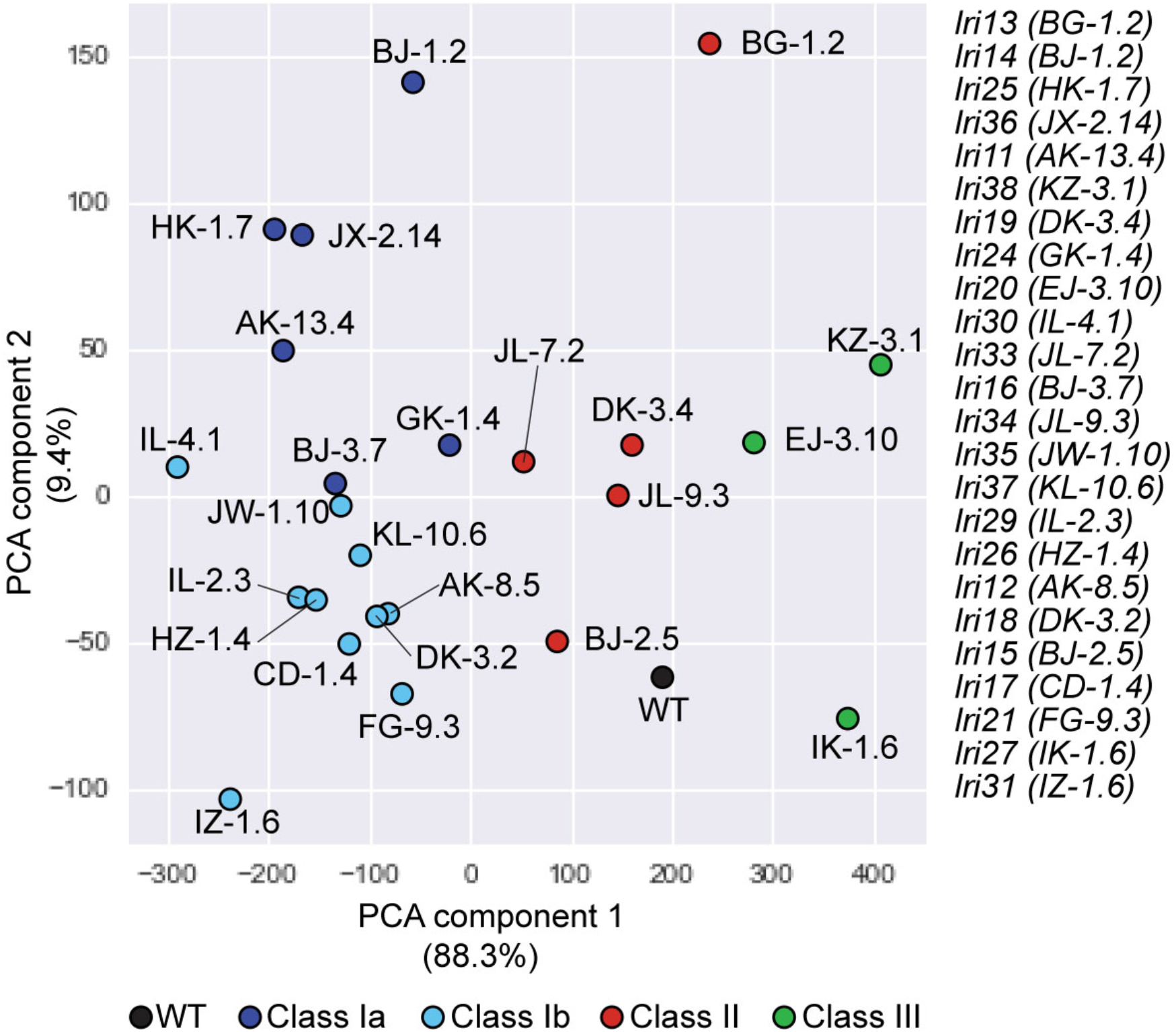
Principal component analysis (PCA) dimensionality reduction analysis-based mutant classification. The PCA dimensionality reduction plot of wild type and recovered mutants based on structural sub-parameters indicates the similarity of phenotypes. Colors denote the defined class of that specimen group: Wild-type (Black), Class Ia (Dark Blue), Class Ib (Light Blue), Class II (Red), and Class III (Green). Proximity of the data points indicates the relative similarity of the recovered mutants; the closer the points, the more closely related the phenotypes are.

### Combination of different node sub-types constructs the intrahepatic biliary network

We previously reported that in wild-type larvae at 5 dpf, the distribution ratios of 3-way, 4-way, and 5-way nodes in the intrahepatic biliary network are relatively constant (16). Here, we further analyzed node types in wild-type larvae at 5 dpf by subdividing into node sub-types: 3-way, 4-way, and 5-way nodes can be subdivided into 3, 4, and 5 node subtypes, respectively (Fig. 4A and B) (Table 2). For instance, there are 3 sub-types of 3-way nodes: a 3-way node connecting to three other nodes (3W3N0E), a 3-way node connecting to two other nodes and one endpoint (3W2N1E), and a 3-way node connecting to one other node and two endpoints (3W1N2E). In wild-type larvae at 5 dpf, these node subtypes show a typical distribution ratio in which the 3W2N1E node ratio (35.74% ± 8.18 s.d.) is always higher than that of the 3W1N2E node (0.93% ± 3.02 s.d.) (Fig. 4C). Similarly, among the 4-way nodes, the ratio of the 4W3N1E (7.60% ± 2.30 s.d.) node is always higher than that of the 4W1N3E (0.97% ± 0.92 s.d.) node (Fig. 4D). The similar pattern is observed in 5-way node subtypes (Fig. 4E). In wild-type larvae, the zebrafish intrahepatic biliary network branching patterns are not identical from one larva to another; however, these data together indicate that there are certain rules and ranges that govern the composition of node-subtypes constructing the intrahepatic biliary network in wild-type larvae.

**Figure 4.**
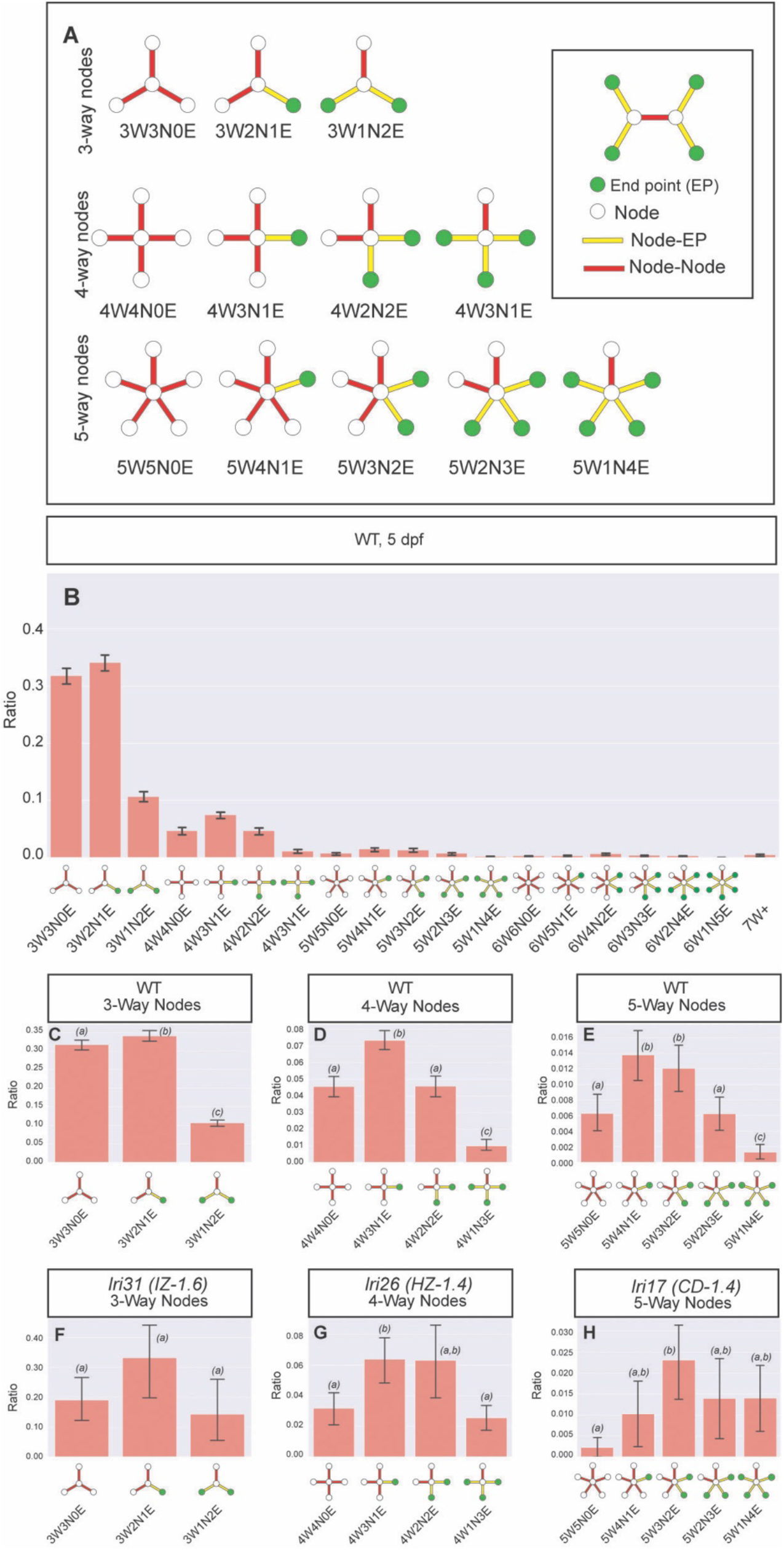
Defining the node-type composition of the intrahepatic biliary network. Computational network analysis revealed that the intrahepatic biliary network could be defined as a combination of different types of nodes. **(A)** Schematics of node-subtypes existing in the intrahepatic biliary network. **(B)** Ratio of node sub-type distribution in the intrahepatic biliary network in wild-type larvae at 5 dpf. X-axis labels denote the number of branches connected to the node (#W), how many of those branches are node-node connections (#N), and how many of those branches are node-endpoint connections (#E). In the schematics, node (white circle), endpoint (green circle), node-node (red bar) and node-endpoint (yellow bar) connections indicate each node sub-type compositions. **(C)** Ratio of 3-way node sub-types in wild-type (WT) is plotted separately. The 3W2N1E node always exists significantly higher than the 3W1N2E node in WT. **(D)** Ratio of 4-way node sub-types in WT is plotted separately. Similar to the distribution of 3-way nodes, the 4W3N1E node always exists with the highest ratio in WT while the 4W1N3E node is always lower than other node sub-types. **(E)** Ratio of 5-way node sub-types in WT is plotted separately. The 5W1N4E node always exists at the lowest ratio in WT. **(F)** Ratio of 3-way node sub-types in *lri31 (IZ-1.6)* mutant larvae at 5 dpf. The ratio difference between 3W2N1E and 3W1N2E is lost in *lri31* mutant larvae. **(G)** Ratio of 4-way node sub-types in *lri26 (HZ-1.4)* mutant larvae at 5 dpf. The relative ratio of the 4W1N3E node was increased in *lri26* mutant larvae. **(H)** Ratio of 5-way node sub-types in *lri17 (CD-1.4)* mutant larvae at 5 dpf. The relative ratio of the 5W1N4E node is increased in *lri17* mutant larvae. These data indicate that that the distribution of node sub-types is regulated in part by these mutations. Bars with a shared letter indicate that the difference is not statistically significant.

**Table 2.** Node sub-type distribution in wild-type larvae at 5 dpf.

| **Node Sub-Type** | **Average** | **Standard Deviation** |
| --- | --- | --- |
| 3W3N0E | 31.45% | 5.18 |
| 3W2N1E | 35.74% | 8.18 |
| 3W1N2E | 10.93% | 3.02 |
| 4W4N0E | 4.62% | 1.92 |
| 4W3N1E | 7.60% | 2.30 |
| 4W2N2E | 4.81% | 2.17 |
| 4W1N3E | 0.97% | 0.92 |
| 5W5N0E | 0.57% | 0.68 |
| 5W4N1E | 1.49% | 1.03 |
| 5W3N2E | 1.37% | 1.07 |
| 5W2N3E | 0.75% | 0.69 |
| 5W1N4E | 0.16% | 0.32 |
| 6W6N0E | 0.15% | 0.27 |
| 6W5N1E | 0.23% | 0.31 |
| 6W4N2E | 0.51% | 0.59 |
| 6W3N3E | 0.24% | 0.37 |
| 6W2N4E | 0.13% | 0.26 |
| 6W1N5E | 0.00% | 0.00 |

Next, we investigated node sub-type distributions in all recovered mutants to find which mutation has the strongest effect on these distributions (Supplementary Fig. 1). Although each of the many mutations has a unique effect on the node sub-type distribution, we found that among the 24 mutants, the *lri31* mutation has the strongest effect on 3-way node sub-type distribution. In *lri31* mutant larvae, the ratio of the 3W2N1E (14.26% ± 13.68 s.d.) node is no longer significantly different from that of the 3W1N2E node (33.16% ± 14.84 s.d.) (Fig. 4F), suggesting that the *lri31* gene influences the 3-way node sub-type distribution. Similarly, among the recovered mutants, 4-way node subtype distribution is most affected in *lri26* mutant larvae, in which the ratio of 4W4N0E (3.16% ± 1.77 s.d.) is no longer significantly higher than that of 4W1N3E (2.55% ± 1.41 s.d.) (Fig. 4G). 5-way node subtype distribution is most affected in *lri17* mutant larvae, in which the ratio of 5W4N1E (1.03% ± 1.17 s.d.) is no longer significantly higher than that of 5W1N4E (1.40% ± 1.15 s.d.) (Fig. 4H). Together, these data suggest that we have identified mutations that influence the proper distribution of node sub-types constructing the intrahepatic biliary network.

### Mutations that affect connection properties

The computational network structure analysis calculates the thickness and length of all individual connections within the intrahepatic biliary network. We used the kernel density estimation algorithm to visualize the mean bivariate distribution of all connection properties in wild-type larvae at 5 dpf (Fig. 5A). We found that all wild-type larvae examined show typical connection property distribution patterns in the plot (Fig. 5A), indicating that all node-node and node-endpoint connections composing the wild-type intrahepatic biliary network have a defined range of thickness and length at this stage. The distribution range of node-node connections is wider for both thickness and length than those of node-endpoint connections (Fig. 5A’ and A’’). The peak distribution of node-node connection properties is always slightly longer in length and wider in thickness than those in node-endpoint connections (Fig. 5A). These data together indicate that in wild-type larvae, connection length and thickness properties show relatively constant distributions, and there must be mechanisms to regulate these distributions.

**Figure 5.**
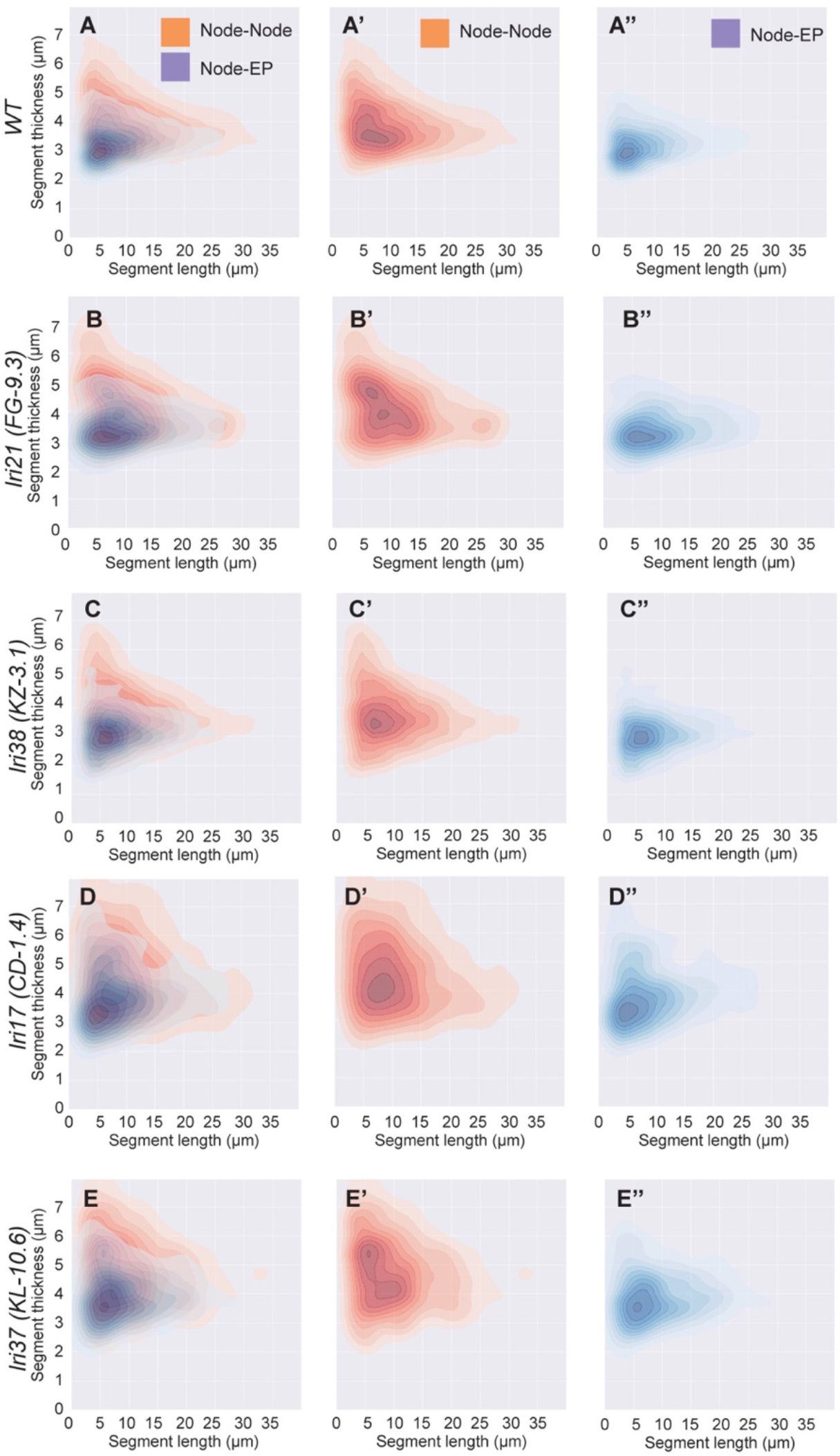
Kernel density estimation based bivariate distribution of connection properties. **(A)** Kernel density estimation plot of thickness and length of node-node (Red) and node-endpoint (Blue) connections in wild-type at 5 dpf. Node-node (Red) and node-endpoint (Blue) plots are shown separately in (A’) and (A’’), respectively. Both node-node and node-endpoint distributions show typical patterns in wild-type larvae. **(B-E)** Kernel density estimation plot of thickness and length of node-node (Red) and node-endpoint (Blue) connections in *lri21 (FG-9.3*) (B), *lri38* (*KZ-3.1*) (C), *lri17* (*CD-1.4*), (D), and *lri37* (*KL-10.6*) mutant (E) mutant larvae at 5dpf. Node-node (Red) and node-endpoint (Blue) plots are shown separately in (B’-E’) and (B’’-E’’), respectively. These mutations uniquely affect node-node and/or node-endpoint segment distributions and make the distribution significantly different from those of wild-type, suggesting that the typical distribution pattern in wild-type is regulated in part by these genes.

To understand which mutations can affect the connection property distributions, we next made kernel density estimation bivariate distribution plots of all recovered mutants (Supplementary Fig. 2). We found that in *lri21* mutant larvae, node-node connection properties are changed significantly from that of wild-type, while node-endpoint connection properties are relatively unchanged (Fig. 5B), suggesting that node-node connections are selectively affected by the *lri21* mutation (Fig. 5B). In contrast, in *lri38* mutant larvae, node-endpoint connection properties became slightly shorter and thicker than those of wild-type (Fig. 5C), suggesting that this mutation affects node-endpoint projections. In *lri17* mutant larvae, the distributions of both node-node and node-endpoint connection properties are significantly different from those in wild-type (Fig. 5D), suggesting that the *lri17* mutation affects all connection properties. In *lri37* mutant larvae, the peak of the node-node connection value distributions moved away from the peak of those in the node-endpoint connection (Fig. 5E). Together, these data suggest that we have identified mutations that affect different subsets of connection properties in the intrahepatic biliary network.

### Network structural heat map based characterization of Class III mutant phenotypes

The connection property distribution analysis revealed that in mutant larvae belonging to Class III, *lri20*, *lri27*, and *lri38*, node-node connections become thicker than those in wild-type (Supplementary Fig. 2). However, where in the liver these thicker node-node connections are localized is not clear. To test the hypothesis that property altered connections could be distributed in a particular area of the liver in these mutants, we have developed additional visualization programs. We have previously generated an algorithm to make two-color network heat maps indicating network crowdedness (16). We modified the procedure (Materials and Methods) to generate a seven-color network heat map of the intrahepatic biliary network in wild-type larvae that shows the density of the network (Fig. 6A). Since recovered mutants belonging to Class III show a phenotype that increases network density, we first generated seven-colored network density heat maps for *lri20*, *lri27* and *lri38* mutant larvae at 5 dpf (Fig. 6B–D). Consistent with the initial analyses, these mutants show increased network density; however, the network density appears to become much higher in a particular area in the liver when compared to the remainder of the liver (Fig. 6B–D), suggesting that network density increases locally inside the liver.

**Figure 6.**
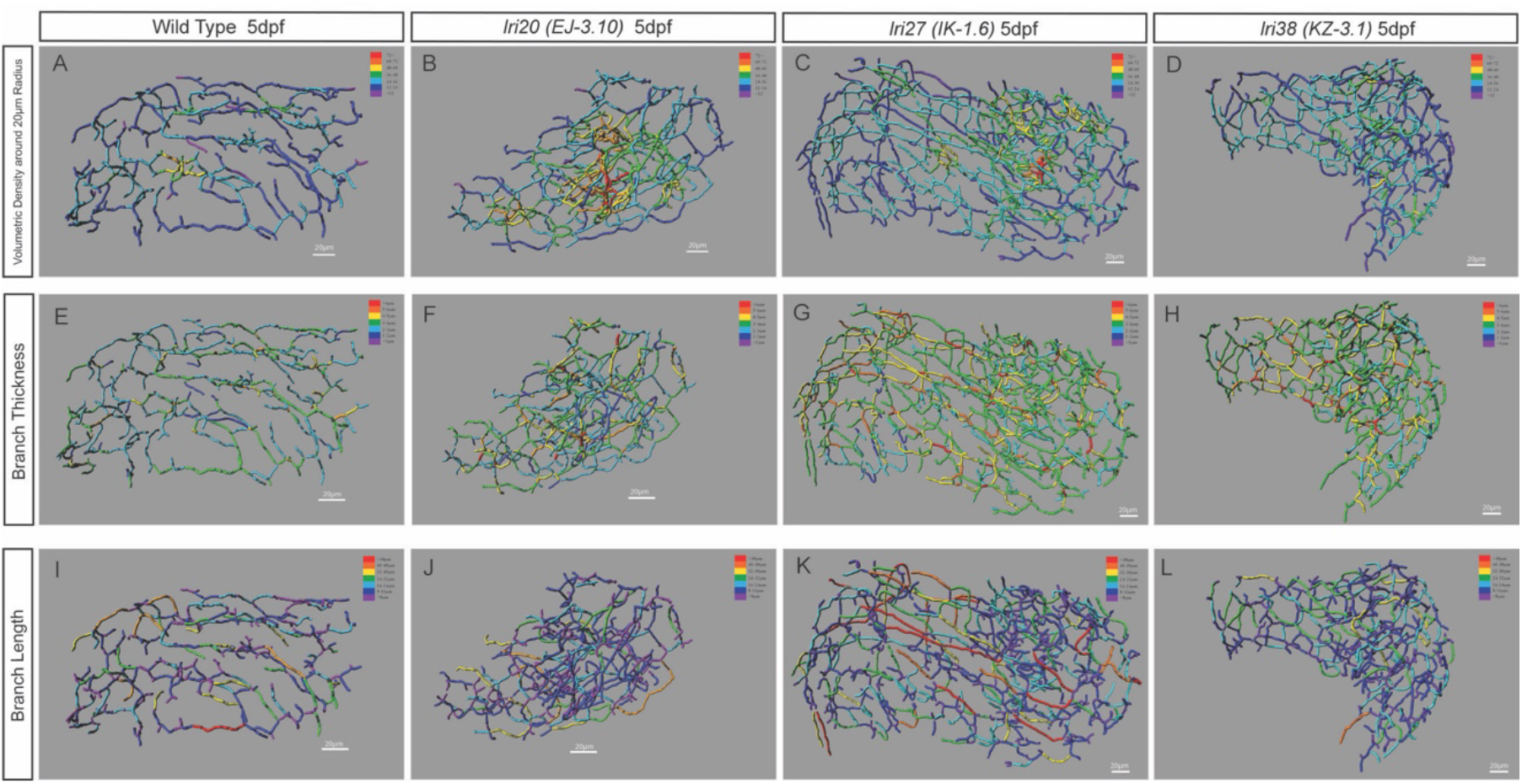
Heat map-based characterization of Class III mutant phenotypes. Heat map skeletal representation of the intrahepatic biliary network showing different segment properties in wild-type (WT) and recovered mutant larvae belonging to Class III. The colors of the heat maps indicate value ranges for each segment property. The hotter the color, the higher the value for all heat maps. **(A-D)** Network volumetric density heat map in WT (A), *lri20 (EJ-3.10)* (B), *lri27 (IK-1.6)* (C) and *lri38 (KZ-3.1)* (D) mutant larvae at 5 dpf. Purple represents less than 12 A.U., dark blue represents 12-24 A.U., light blue represents 24-36 A.U., green represents 36-48 A.U., yellow represents 48-60 A.U., orange represents 60-72 A.U., and red represents 72 A.U. and greater. These mutants show increased network density in a particular area in the liver that is not seen in WT. **(E-H)** Network connection thickness heat map in WT (E), *lri20 (EJ-3.10)* (F), *lri27 (IK-1.6)* (G) and *lri38 (KZ-3.1)* (H) mutant larvae at 5 dpf. Purple represents less than 1 μm, dark blue represents 1-2 μm, light blue represents 2-3 μm, green represents 3-4 μm, yellow represents 4-5 μm, orange represents 5-6 μm, and red represents 6 μm and greater. Network thickness increased uniformly across the entire network in the mutants compared to WT. **(I-L)** Network connection length heat map in WT (I), *lri20 (EJ-3.10)* (J), *lri27 (IK-1.6)* (K) and *lri38 (KZ-3.1)* (L) mutant larvae at 5 dpf. Purple represents less than 8 μm, dark blue represents 8-16 μm, light blue represents 16-24 μm, green represents 24-32 μm, yellow represents 32-40 μm, orange represents 40-48 μm, and red represents 48 μm and greater. Distribution of longer connections was different from those in WT larvae only in *lri27* mutant larvae.

We then hypothesized that network density might increase where segments are thicker in mutants belonging to Class III. However, when we generated network heat maps indicating connection thickness in *lri20*, *lri27* and *lri38* mutant larvae (Fig. 6F–G), we found that network thickness was increased uniformly across the entire network in these mutants compared to that of wild-type larvae (Fig. 6E). Thus, we concluded that connection thickness and density are determined independently in these mutants. We also generated network heat maps indicating connection segment length in *lri20*, *lri27* and *lri38* mutant larvae (Fig. 6J–L), and we found that distribution of longer connections appeared to be different from those in wild-type larvae only in *lri27* mutant larvae (Fig. 6I). Together, these data highlight the usefulness of generating network heat maps to understand where significant property changes develop in the liver.

### Mutations that affect the integrity of the intrahepatic biliary network

Among 24 recovered mutants, we found that 4 mutants belonging to Class I show the additional phenotype in which some *Tg(Tp1-MmHbb:EGFP)^um14^*-expressing biliary epithelial cells (BECs) segregate from the intrahepatic biliary network. Here, we referred to this phenotype as the BEC segregation phenotype (Table 1). For instance, in *lri31* mutant larvae at 5 dpf, single *Tg(Tp1-MmHbb:EGFP)^um14^*-expressing BECs located in the anterior part of the liver segregate from the continuous intrahepatic biliary network (Fig. 7B), while all *Tg(Tp1-MmHbb:EGFP)^um14^*-expressing BECs are always connected to the one continuous intrahepatic biliary network in wild-type larvae (Fig. 7A). The segregated single BECs frequently make direct contact with the *Tg(kdrl:RFP_CAAX)^y171^*–expressing intrahepatic vascular network (Fig. 7B and C). These data indicate that these four genes are responsible for maintaining the integrity of the continuous network.

**Figure 7.**
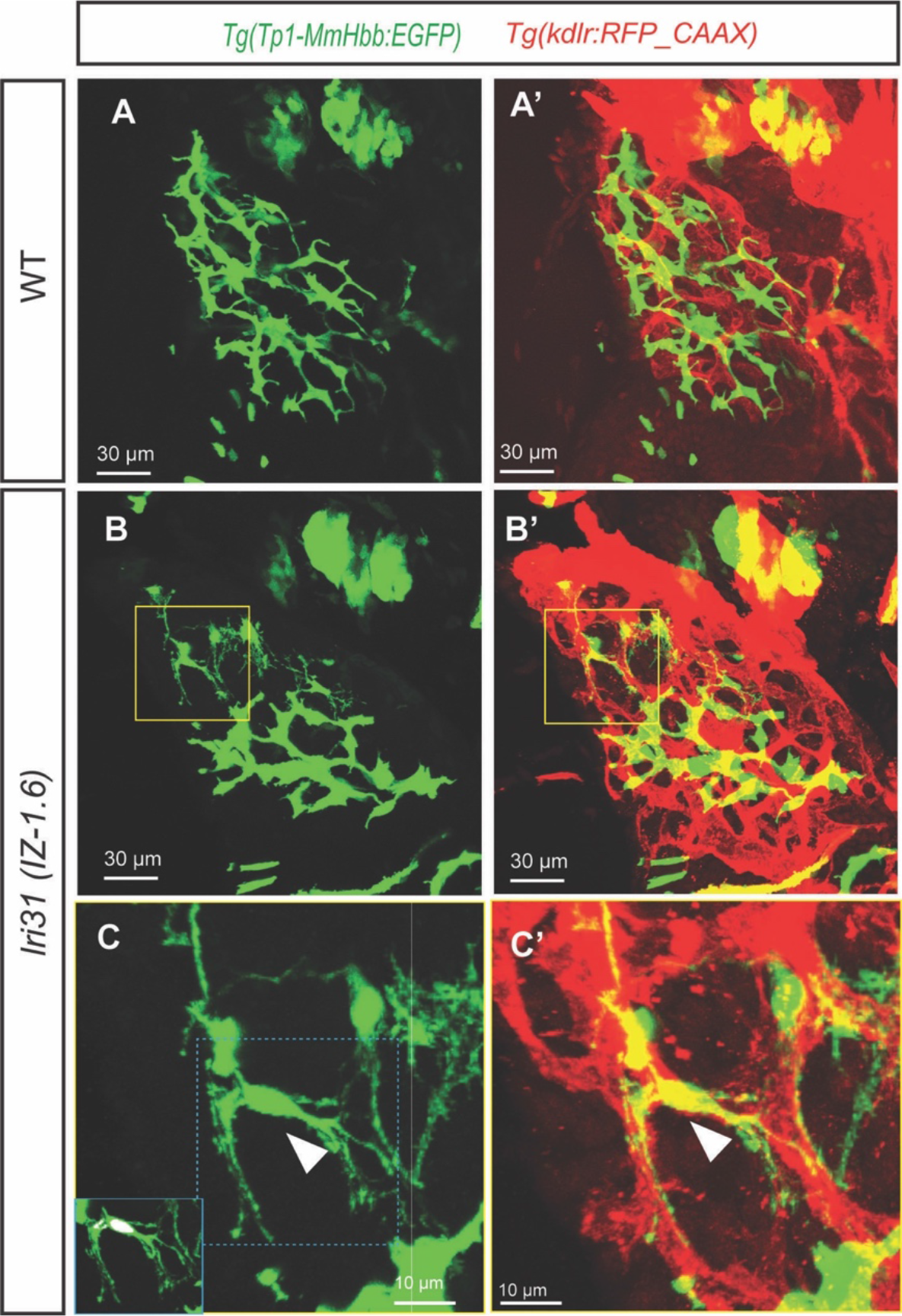
Biliary epithelial cells segregate from the intrahepatic biliary network in *lri31 (IZ-1.6)* mutant larvae at 5 dpf. **(A-C)** Projected confocal images of wild-type (A) and *lri31 (IZ-1.6)* mutant (B and C) larvae visualized for *Tg(Tp1-MmHbb:EGFP)^um14^* expressions at 5 dpf. *Tg(kdrl:RFP_CAAX)^y171^* expressions are merged in (A’-C’). Yellow-outlined area in B is magnified and shown in C. Blue-outlined area in C is magnified and merged with the DAPI and GFP co-localization signal in the left bottom corner. Ventral views, anterior to the top. In *lri31 (IZ-1.6)* mutant larvae, single biliary epithelial cells segregate from the intrahepatic biliary network and make physical contact with the intrahepatic vascular network (White arrowhead in C).

### Some recovered mutants show biliary system phenotypes in the adult liver

Likely because we screened for mutants that show no gross morphological phenotype, the majority of mutations we collected in this screen are viable (19 out of 24) (Table 1). We hypothesized that these viable mutations might continue to affect the intrahepatic biliary network in the adult stage. To test this hypothesis, we collected recovered mutant larvae at 5 dpf and raised homozygous mutant fish into the adult stage. We then examined *Tg(Tp1-MmHbb:EGFP)^um14^* expression in the adult liver (6 months post-fertilization) of these homozygous mutant fish (Fig. 8). In the wild-type adult liver, *Tg(Tp1-MmHbb:EGFP)^um14^* remains to be expressed in the intrahepatic biliary network (Fig. 5A). In the liver of *lri37* mutant fish, *Tg(Tp1-MmHbb:EGFP)^um14^* expression in the liver appears similar to that of wild-type (Fig. 8B), suggesting that the intrahepatic biliary network is impaired in *lri37* mutant larvae but recovers in adult fish. In the *lri26* (Fig. 8C) and *lri14* (Fig. 8D) mutant adult livers, *Tg(Tp1-MmHbb:EGFP)^um14^* expression in the liver is slightly reduced compared to that of wild-type fish, suggesting that the intrahepatic biliary network in these mutant fish is reduced. In the *lri13* (Fig. 8E), *lri21* (Fig. 8F), and *lri24* (Fig. 8G) mutant adult livers, *Tg(Tp1-MmHbb:EGFP)^um14^* expression in the liver is greatly reduced and nearly missing, suggesting that the intrahepatic biliary network is almost lost in these mutants. In *lri29* (Fig. 8I) and *lri20* (Fig. 8J) mutant adult livers, the *Tg(Tp1-MmHbb:EGFP)^um14^*-expressing intrahepatic biliary network becomes thicker compared to that of the wild-type adult liver, suggesting that the failure of BEC segregation might cause thicker networks to form in these mutant fish. Together, these data indicate that we have established new zebrafish models for adult biliary system defects, which can potentially be used as cholestatic liver disease animal models.

**Figure 8.**
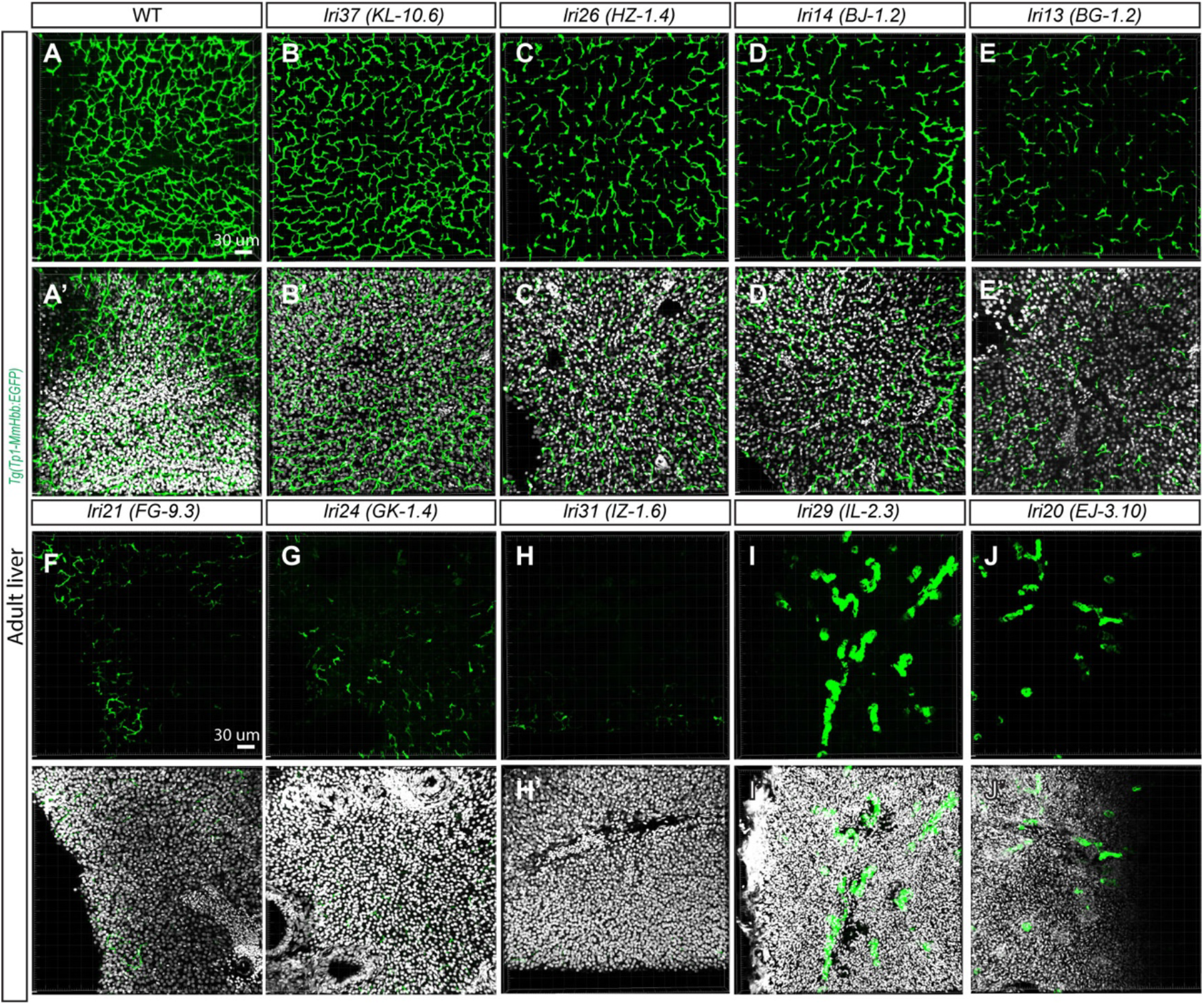
Projected confocal images of wild-type and mutant adult livers. **(A-J)** Projected confocal images of the adult liver visualized for *Tg(Tp1-MmHbb:EGFP)^um14^* expression in wild-type (A), *lri37 (KL-10.6)* (B), *lri26 (HZ-1.4)* (C), *lri14 (BJ-1.2)* (D), *lri13 (BG-1.2)* (E), *lri21 (FG-9.3)* (F), *lri24 (GK-1.4)* (G), *lri31 (IZ-1.6)* (H), *lri29 (IL-2.3)* (I), and *lri20 (EJ-3.10)* (J) mutant fish. DAPI staining is overlaid in (A’-J’). The cross sections of the liver were used.

## Discussion

In this study from a forward genetic screen, we have recovered 24 new mutants which show specific defects in the branching patterns of the intrahepatic biliary network. This screen was successful only because we utilized the previously developed network structure analysis algorithm to confirm the subtle difference in the branching patterns. By the algorithm, we converted the phenotypic difference in these 24 mutants into multi-dimensional arrays and applied machine-learning-based unsupervised classification approaches. These approaches objectively classified these 24 mutants into three major classes and quantified phenotypical similarities among them.

The clustered heat map displays a large amount of data in an intuitive format that facilitates the detection of hidden structures and relations in the dataset. We classified recovered mutant phenotypes based on this algorithm. The key question to be evaluated is the extent to which unsupervised clustering reflects the signaling pathways regulated by the mutated genes. Until recently, mapping of a causative mutation of ENU mutagenesis-derived mutants was an extremely labor-intensive process. However, the development of next-generation sequencing-based approaches (19–22) has made this process significantly easier. In fact, we have already identified that the *lri35* mutation disrupts the *nckap1l* gene by solely using decent size (total 22 millions reads) RNAseq data (23). We initially focused on the *lri35* mutation because its phenotype is the most similar to that of Cdk-5 suppressed larvae based on the Euclidean distance of the network sub-parameters (23). We then demonstrated that the *lri35* mutation genetically interacts with the Cdk5 pathway (23), highlighting that precise phenotype quantification can predict the signaling pathway that the mutation disrupts. Further identifying the responsible genes mutated by the rest of the 23 recovered mutations will clearly verify the accuracy of the unsupervised classification achieved in this study.

Genetic complementation tests within the same class have been completed and none of the tested pairs within the same class complemented one another, suggesting that these 24 mutations affect 24 different genes. It should be noted that, during the complementation test, there were a few cases in which less than five percent of offspring showed much milder phenotypes than those seen in their parents. Since all recovered mutations are full penetrant and recessive, we concluded these results could be due to genetic interaction between two loci. The number of mutants we recovered from this screening is significantly higher than the number of genes already known to be involved in zebrafish biliary system formation. Thus, identifying the responsible genes mutated by these mutations will reveal previously unappreciated genes playing essential roles in biliary system formation. The complementation test results also indicated that a genetic screen focusing on the biliary system has not been saturated; more mutations remain to be discovered. However, different alleles on the same gene could potentially produce very distinct phenotypes which could be classified into different classes. Thus, it is still possible that some recovered mutations could affect the same gene.

In this study, we redefined node subtypes existing in the wild-type intrahepatic biliary network (Fig. 4). This analysis provided a previously undescribed view on how the network is constructed. We found that two node subtypes, 3W3N0E and 3W2N1E, compose more than 65% of the total nodes in the wild-type intrahepatic biliary network at 5 dpf (Fig. 4A). Other node subtype distributions are relatively constant among wild-type larvae at 5 dpf (Fig. 4A), and we found several mutants, including *lri31*, that influence the patterned distributions of node subtypes (Fig. 4, Supplementary Fig. 1). Identifying the responsible genes affected by these mutations will further address molecular pathways responsible for node subtype determination and distribution.

We showed that mutant larvae belonging to Class III display increased network density of the intrahepatic biliary network at 5 dpf (Fig. 6). We previously reported that pharmacological suppression of Pak1 or Lim kinase also induced a phenotype (16) similar to those mutant larvae belonging to Class III; thus, it could be possible that some of the mutants belonging to this class might affect the Pak1 or Lim kinase-mediated signaling pathways.

In this study, we found several mutants in which single biliary epithelial cells segregate from the network (Fig. 7). We only used the remaining continuous biliary network for the skeletal structural analysis, as the segregated single cells are not continuous with the network.

Our high-stringency screening criteria led us to collect viable mutations that will allow us to generate double- and triple-mutant fish. It will be intriguing to examine the biliary network phenotypes in double- and triple-mutant larvae of the same class to examine potential genetic interactions. These viable mutants showing adult liver phenotypes are also an important resource for understanding cholestatic liver disease. A community-based study suggested that cholestasis may affect as much as 10 to 20% of the population (24), while in the majority of the cases, cholestasis remains benign (25). To our surprise, some adult mutant fish lost almost all *Tg(Tp1-MmHbb:EGFP)^um14^*-expressing biliary epithelial cells in the liver (Fig. 8), but remain viable and fertile. These data do not exclude the possibility that in these severe phenotypes, transgenic expression may be lost while the actual biliary epithelial cells persist. In human cholestatic liver disease patients, the obstruction of the intrahepatic biliary network frequently leads to hepatic inflammation; thus, it will be important to examine the degree of hepatic inflammation in these mutant fish. These new viable mutations are an important biological resource for modelling different aspects of human cholestatic liver diseases. We also found that biliary network phenotypes in larvae do not always correlate with biliary network phenotypes in the adult liver. For instance, *lri14* mutant larvae show a reduced complexity phenotype in the biliary network (Class Ia), whereas *lri13* mutant larvae show a crowded biliary network phenotype (Class III). However, these two mutant fish show very similar biliary network phenotypes in the adult liver (Fig. 8; Table 1). These data suggest that in many cases, the mutations that affect larval biliary network formation also influence the intrahepatic biliary network in adults, but the detailed biliary phenotypes in adults need to be examined independently from those of larvae.

Overall, this study combined a zebrafish forward genetic screen with unsupervised classifications to identify 24 new mutations which affect different aspects of biliary network formation and maintenance. These mutations will be a tremendous resource to further understand the molecular pathways governing biliary system formation and molecular pathologies leading to cholestatic liver diseases.

## Materials and Methods

### Zebrafish husbandry and transgenic lines

Zebrafish (*Danio rerio*) larvae were obtained from natural crosses of the wild-type AB/TL strain or heterozygous mutant fish. The following transgenic lines were used: *Tg(Tp1-MmHbb:EGFP)^um14^ (15)* and *Tg(kdrl:RFP_CAAX)^y171^*(26). We also used 24 new recovered mutants (Table 1), which are generated in this study. Animal husbandry methods were approved by the Cleveland Clinic’s Institutional Animal Care and Use Committee.

### Mutant recovery from ENU-based mutagenesis

The standard three-generation screen was conducted in our lab as previously described (17, 18). In brief, total 228 F2 families representing 332.32 genomes were screened for altered *Tg(Tp1-MmHbb:EGFP)^um14^* expression in the liver at 5 dpf, and 24 mutants were recovered and established. The recovered mutants were backcrossed to the original AB/TL strain for five or more generations before starting detailed phenotype analyses.

### Imaging and computational network structure analysis

Confocal z-stack data of *Tg(Tp1-MmHbb:EGFP)^um14^* expression in the liver were obtained using a Leica SP5 confocal microscope as previously described (27). The z-step used on the images was 0.42 μm. We used Imaris 8.2 software (Bitplane) to digitally crop the image such that only EGFP expression from the intrahepatic biliary network remained for further analysis. The Surface feature was used to isolate the biliary network, and the background subtraction threshold adjusted until the network was continuous. The Liver Analysis Program 5.3 (16) was used for all computational network structure analyses. For all analyses, we confirmed the proper skeletal conversion by overlaying the original confocal image with the converted skeletal image as previously described (16).

### Clustered heat map-based classification and PCA analysis

Structural sub-parameters of the intrahepatic biliary network in recovered mutant larvae were calculated by the Liver Analysis Program 5.3 as described above. Each sub-parameter value was normalized to the mean value in wild-type larvae. The clustering hierarchical heat-map (12) was generated based on the normalized mean values of recovered mutant larvae. Mean values of structural sub-parameters of each mutant larva were used to generate a PCA plot (12). The number of samples used per mutant was described in Table 1.

### Heat map analysis on connection segment properties

The data output of the Liver Analysis Program 5.3 was used for further data analysis of the biliary network with the Liver Density Analysis Program v1.4 and Liver Segmentation Selection Program v1.0. The Liver Density Analysis Program v1.4 (16) was used for all computational network density analysis. The Liver Segmentation Selection Program v1.0 was used to isolate segmentation groups for creating heat maps. To create a heat map, the data output of the Liver Analysis Program 5.3 was segmented based on the connection properties. The program output a new TIF stack containing only the segments and nodes of interest. This process was repeated for each class interval. The TIF stacks were then added to Imaris as new channels and colored to denote the class interval. Through this method, heat maps were created for branch length, branch thickness, and network density of the biliary network.

### Refined node-type composition calculation

Based on the initial output of the Liver Analysis Program 5.3, we calculated the frequency of node and endpoint appearance per each existing node to determine node sub-types. This calculation script is named as the Node Sub-type Calculation Program v1.1.

### Kernel density estimation based bivariate distribution plots

We averaged the distribution of the length and thickness in each mutant and plotted the kernel density estimation (28) based bivariate distribution. The averaging was based on the estimation that the distribution in each mutant is relatively similar.

### Adult liver tissue processing and imaging

Adult fish were dissected to examine altered *Tg(Tp1-MmHbb:*EGFP*)^um14^* expression in the intrahepatic biliary network in the adult liver at 6 months post-fertilization. Once euthanized, the fish were dissected in 1x-D-PBS, the liver and intestines removed, and the liver separated into lobes. The lobe was fixed with 2% Formaldehyde in PEM, washed twice with PBS and then embedded in 2% GeneMate LowMelt Agarose (Cat. No. E-3126-25) in PBS. The agarose block was sliced at a thickness of 250 μm with Leica VT1000S vibratome. The slices were then stained overnight with Invitrogen DAPI (Cat. No. D1306). *Tg(Tp1-MmHbb:EGFP)^um14^* expression in the liver was scanned in a z-stack image on a Leica SP5 confocal microscope, and the image was exported from Bitplane Imaris software.

### Statistics

To compare three or more means, one-way ANOVA followed by Tukey’s HSD test was used.

## Author Contribution

T.F.S conceived the research plan. M.D., J.L.L., L.X.P., A.M., G.N., I.G., and T.F.S. conducted a forward genetic screen along with people acknowledged. J.L. analyzed the adult liver phenotypes. K.M.T, K.L.O., M.D., J.L.L., L.X.P., A.M., G.N., and I.G. carried out the confocal observations of mutant larvae. K.L.O., K.T., W.L., K.M.T., M.A.M., and T.F.S. developed and performed computational analyses. D.J.S., K.M.T, K.L.O., and T.F.S. analyzed the data and drafted the manuscript. All authors discussed the results and contributed to the final manuscript.

## Acknowledgements

We thank Madeline Schaub, James Cantrell, Cassandra Bilogan, Kasey Kisewetter, and Kimia Ghaffari for helping our forward genetic screen. We thank Saswat Sahoo for the critical reading of the manuscript. We also thank Maria Roufaeil and Katherine Bell for helping computational analyses.

