## Supplementary Information for "Forward genetics combined with unsupervised classifications identified zebrafish mutants affecting biliary system formation"

**Supplementary Figure 1. Node-type composition of the intrahepatic biliary network in recovered mutant larvae. (A)** Ratio of node sub-type distribution in the intrahepatic biliary network in wild-type larvae at 5 dpf. X-axis labels denote the number of branches connected to the node (#W), how many of those branches are node-node connections (#N), and how many of those branches are node-endpoint connections (#E). **(B-Y)** Ratio of node sub-type distribution in the intrahepatic biliary network in *lri24* (*GK-1.4*) (B), in *lri14* (*BJ-1.2*) (C), *lri24* (*GK-1.4*) (B), *lri14* (*BJ-1.2*) (C), *lri25* (*HK-1.7*) (D), *lri36 (JX-2.14)* (E), *lri11* (*AK-13.4*) (F), *lri16* (*BJ-3.7*) (G), *lri31* (*IZ-1.6*) (H), *lri21* (*FG-9.3*) (I), *lri30* (*IL-4.1*) (J), *lri29* (*IL-2.3*) (K), *lri17* (*CD-1.4*) (L), *lri18* (*DK-3.2*) (M), *lri26* (*HZ-1.4*) (N), *lri35* (*JW-1.10*) (O), *lri12* (*AK-8.5*) (P), *lri37* (*KL-10.6*) (Q), *lri33* (*JL-7.2*) (R), *lri15* (*BJ-2.5*) (S), *lri34* (*JL-9.3*) (T), *lri13* (*BG-1.2*) (U), *lri19* (*DK-3.4*) (V), *lri20* (*EJ-3.10*) (W), *lri27* (*IK-1.6*) (X), and *lri38* (*KZ-3.1*) (Y) mutant larvae at 5 dpf.

**Supplementary Figure 2. Kernel density estimation based bivariate distribution of connection segment properties in recovered mutant larvae. (A)** Kernel density estimation plot of thickness and length of node-node (Red) and node-endpoint (Blue) connections in wild-type at 5 dpf. Node-node (Red) and node-endpoint (Blue) plots are shown separately in (A') and (A''), respectively. Both node-node and node-endpoint distributions show typical patterns in wild-type larvae. **(B-Y)** Kernel density estimation plot of thickness and length of node-node (Red) and node-endpoint (Blue) connections in *lri24* (*GK-1.4*) (B), *lri14* (*BJ-1.2*) (C), *lri25* (*HK-1.7*) (D), *lri36 (JX-2.14)* (E), *lri11* (*AK-13.4*) (F), *lri16* (*BJ-3.7*) (G), *lri31* (*IZ-1.6*) (H), *lri21* (*FG-9.3*) (I), *lri30* (*IL-4.1*) (J), *lri29* (*IL-2.3*) (K), *lri17* (*CD-1.4*) (L), *lri18* (*DK-3.2*) (M), *lri26* (*HZ-1.4*) (N), *lri35* (*JW-1.10*) (O), *lri12* (*AK-8.5*) (P), *lri37* (*KL-10.6*) (Q), *lri33* (*JL-7.2*) (R), *lri15* (*BJ-2.5*) (S), *lri34* (*JL-9.3*) (T),

*Iri13* (*BG-1.2*) (U), *Iri19* (*DK-3.4*) (V), *Iri20* (*EJ-3.10*) (W), *Iri27* (*IK-1.6*) (X), and *Iri38* (*KZ-3.1*) (Y) mutant larvae at 5dpf. Node-node (Red) and node-endpoint (Blue) plots are shown separately in (B'-Y') and (B''-Y''), respectively. These mutations uniquely affect node-node and/or node-endpoint segment distributions and make the distribution significantly different from those of wild-type, indicating that the typical distribution pattern in wild-type is regulated in part by these genes.**Supplementary Fig. 1**

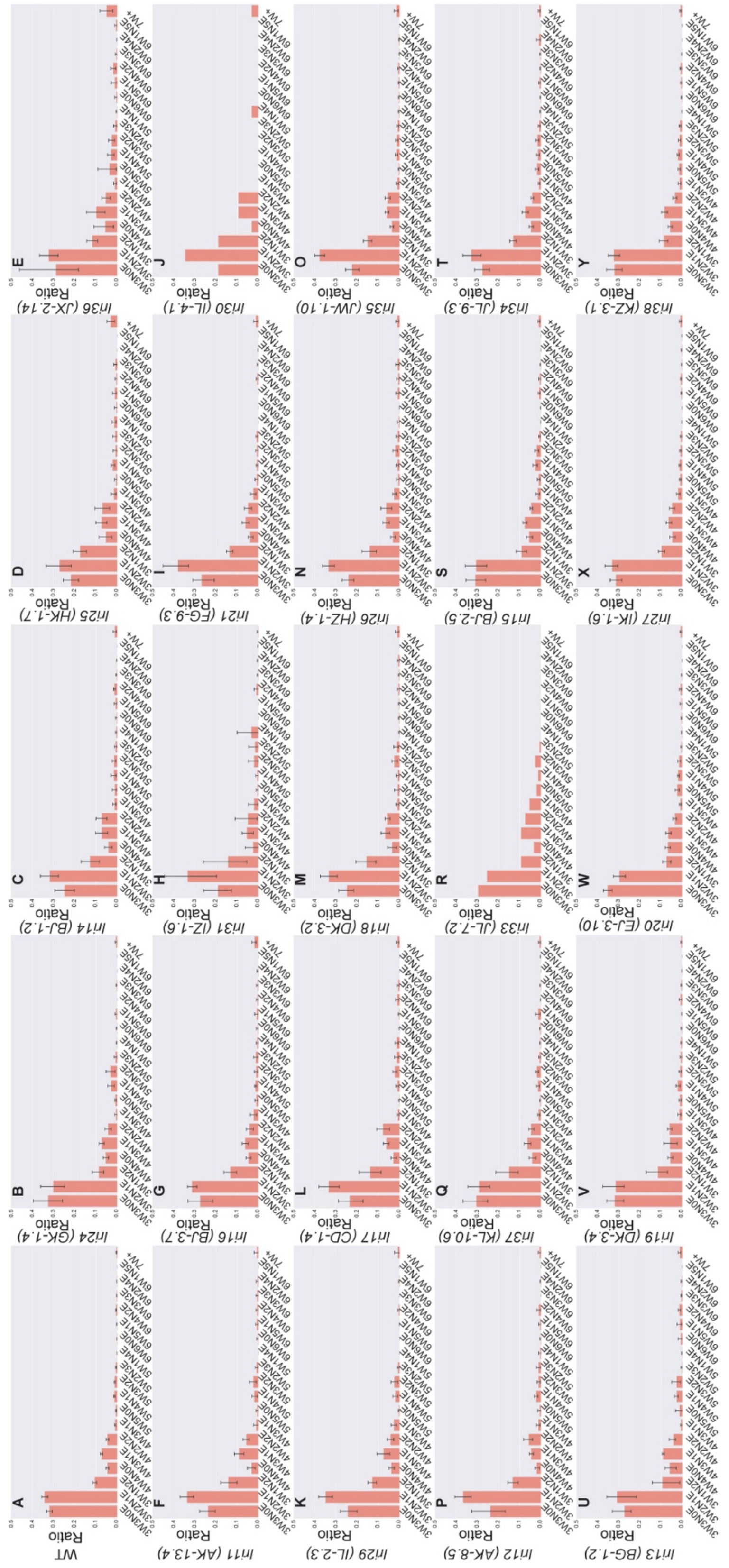
A WT
B In24 (GK-1.4)
C In14 (BJ-1.2)
D In25 (HK-1.7)
E In36 (JX-2.14)
F In11 (AK-13.4)
G In16 (BJ-3.7)
H In31 (IZ-1.6)
I In21 (EG-9.3)
J In30 (IL-4.1)
K In29 (IL-2.3)
L In17 (CD-1.4)
M In18 (DK-3.2)
N In26 (HZ-1.4)
O In35 (JW-1.10)
P In12 (AK-8.5)
Q In37 (KL-10.6)
R In33 (JL-7.2)
S In15 (BJ-2.5)
T In34 (JL-9.3)
U In13 (BG-1.2)
V In19 (DK-3.4)
W In20 (EJ-3.10)
X In27 (IK-1.6)
Y In38 (KZ-3.1)
Ratio

**Supplementary Figure 2.**

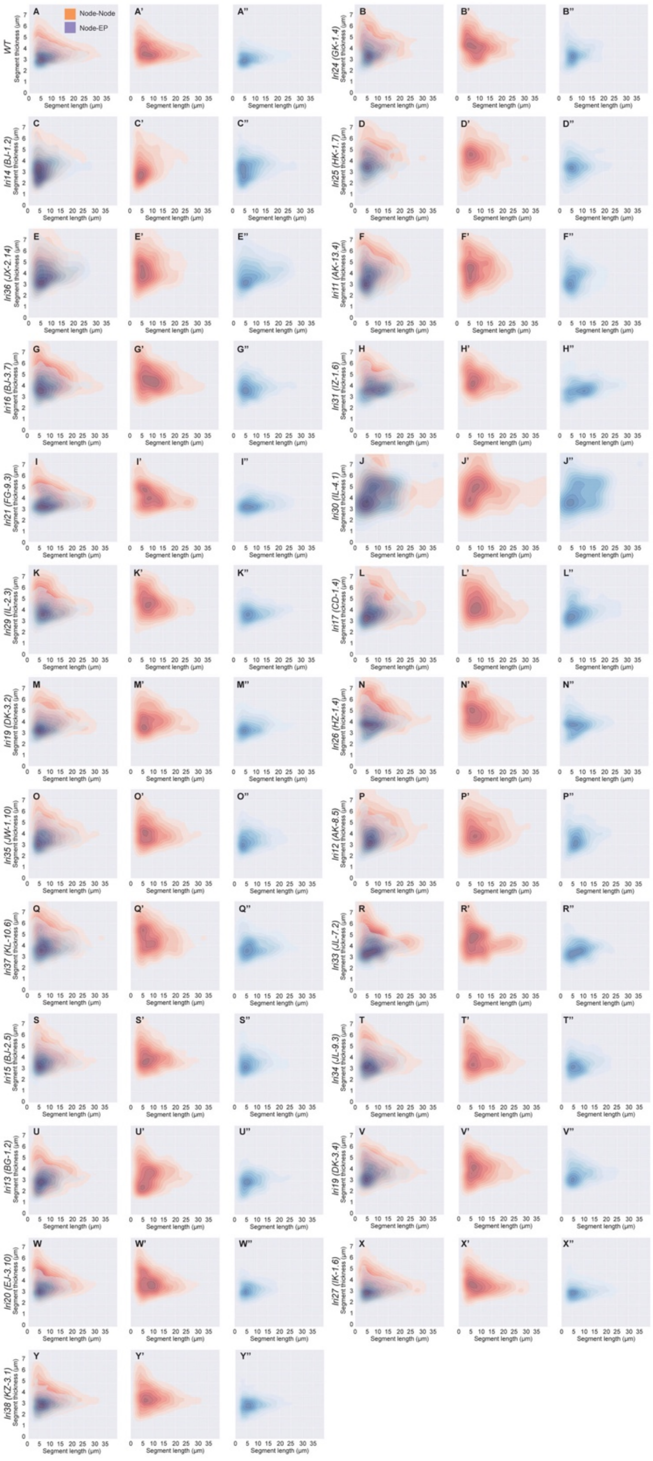
